# Anti-allergic Effect of Soluble Beta-glucan on Japanese Cedar Pollinosis and Mechanism Underlying Its Allergy-ameliorating Action

**DOI:** 10.1101/2025.10.26.684603

**Authors:** Mutsumi Tanaka, Tetsu Sawamoto, Keiko Mizuyama, Fusako Mitsunaga, Shin Nakamura

## Abstract

Japanese cedar (sugi) pollinosis is triggered by sensitization to sugi pollen, which contains sugi antigens, sugi basic proteins (SBP: Cryj1 and Cryj2), and adjuvant polysaccharides, insoluble β-glucan particles (IBG: a dectin-1 agonist). The presence of IBG in sugi pollen enhances the production of SBP-specific IgE, which participates in inflammatory response via activation of mast cells to lead to their degranulation, resulting in sneezing. Another *β*-glucan derived from black yeast, soluble *β*-glucan (SBG: a dectin-1 antagonist), is known to have an anti-allergic function.

Herein, we demonstrated that oral administration of SBG suppressed both SBP-specific IgE production and SBP-induced sneeze in sugi pollinosis mice model. DNA microarray analyses revealed previously unknown mechanisms underlying SBG-mediated anti-allergy action. SBG ameliorated the dysregulated expression of 16 immune-inflammatory genes that were unusually either up- or down-regulated in sugi pollinosis mice. Based on the DNA microarray informatics, it was suggested that SBG mitigated the IBG-mediated hyper-response of IgE production and inflammatory reaction in immune-inflammatory cells, B and T cells, antigen-presenting cells, and granulocytes (neutrophils and eosinophils) within iliac lymph nodes. The SBG-mediated inhibition of SBP-induced sneezing in pollinosis mice results from a decline in SBP-specific IgE level. The oral administration of SBG influenced gut microbiota, notably enhancing beneficial bacteria such as bifidobacteria. Consequently, it is plausible that short-chain fatty acids, produced by these beneficial bacteria, inhibited mast cell degranulation, leading to a reduction in sneezing.

Collectively, SBG demonstrated anti-allergic effects against sugi pollinosis by inhibiting the immune response, specifically SBP-induced IgE production, as well as reducing the inflammatory reaction characterized by SBP-induced sneezing.

## 1. Introduction

*β*-glucans are naturally occurring polysaccharides of D-glucose monomers linked by *β*-glycosidic bonds, which feature a *β*-1,3-glucan backbone with diverse branching structures. *β*-glucans exhibit a well-defined structure–activity relationships; differences across their various sources and structures result in distinct biological activities. The physicochemical characteristics of *β*-glucans can differ, such as some may be insoluble, form aggregates, and have high molecular weight, whereas others may be soluble, do not form aggregates, and have low molecular weight. These differences in physicochemical properties are critical and influence different biological activities, including immunomodulation [1]. As β-glucans are exogenous compounds, such as pathogens, for mammals, including rodents and primates, they are recognized by pattern recognition receptors (PRRs) to evoke intracellular β-glucan signaling machinery [1, 2]. Dectin-1 is a type II C-lectin-like membrane receptor of the PRR family and is expressed on the surface of myeloid cells (including lymphocytes, macrophages, neutrophils, and mast cells). *β*-glucans bind to the carbohydrate recognition domain of dectin-1. Notably, insoluble *β*-glucan particles (IBG) are able to induce oligomerization and elicit dectin-1 signaling, resulting in Src activation and Syk recruiting, but soluble *β*-glucans (SBG) are unable to do this [1,3,4]. IBG stimulates an immune-inflammatory response that alters gene expression, cytokine and chemokine production, antigen presentation to T and B cells, and immunoglobulin production [1]. Conversely, SBG inhibits the stimulative actions of IBG by competitively binding to dectin-1, thereby acting as a dectin-1 antagonist. Pollen allergy is an IgE-mediated type I hypersensitivity reaction in which the production of antigen-specific IgE antibodies and the activation of mast cells are the limiting factors. Inclination toward a type 2 immune state, Th2, promotes the production of IgE antibodies directed toward a specific antigen called an allergen. IgE antibodies bind to high-affinity IgE receptors (FcεRI) on mast cells and basophils, activating them to trigger degranulation for the release of varied inflammatory mediators, including histamine. Pollen is not only an allergen carrier but also an adjuvant during the induction phase of the allergic immune response [5].

Callose is a major component of higher plant pollen and has a polysaccharide nature with *β*-1,3-glucan [6, 7]. Pollen from Japanese cedar, known as sugi, contained dectin-1 reactive *β*-glucan [8], which induces the production of sugi-allergen-specific IgE in mice [9]. The C-type lectin receptor, dectin-1, was known to be involved in the polarization of CD4+ T cells towards specific Th2 responses associated with specific IgE production and allergic diseases [10]. In a previous paper it was reported that sugi pollen contained IBGs, which had a potent adjuvant activity via binding to dectin-1 on both antigen-presenting and lymphoid cells, leading to an allergic response to produce a specific IgE antibody towards sugi allergen, sugi basic protein (SBP), Crj-1 and Crj-2 [11].

SBG is another type of *β*-glucan with *β*-1,6-branched structures and is mainly produced by the black yeast, Aureobasidium pullulans [12]. The SBG obtained from black yeasts are recognized for their immunoregulatory properties, which can alleviate allergic and/or inflammatory disorders [12, 13]. SBGs and their related compounds function as dectin-1 antagonists and are getting attention for use as anti-allergic and anti-inflammatory materials. However, detailed information on the mechanism underlying SBG-mediated anti-allergic action has not been sufficiently accumulated yet.

In this study, we aimed to examine the anti-allergic effects of SBG and the inhibition of both SBP-specific IgE production and SBP-induced sneezing in a mouse model of sugi-pollinosis. We also aimed to elucidate the mechanism underlying the SBG-evoked anti-allergic action using a DNA microarray, yielding previously unknown molecular events that occurred in the iliac lymph nodes of the pollinosis model.

## 2. Materials and Methods

### 2.1. Reagents

The following materials were used in this study: phosphate-buffered saline (PBS) (Nissui, Tokyo, Japan), a Nunc-immunomodule, F8 Maxisorp, streptavidin-horseradish peroxidase (SA-HRP) (Thermo Fisher Scientific K.K., Yokohama, Japan), and tetramethylbenzidine (TMB) (Sigma-Aldrich Co. LLC, Tokyo, Japan). Casein Na, sodium azide (NaN3), and Tween 20 were purchased from FUJIFILM Wako Pure Chemical Corporation (Osaka, Japan). An ELAST enzyme-linked immunosorbent assay (ELISA) amplification system (PerkinElmer, Inc., Waltham, MA, USA), anti-mouse IgE (ab11580; Abcam, Cambridge,UK), streptavidin-horseradish peroxidase conjugate (SA10001; Invitrogen, Carlsbad, CA, USA), and biotynyltyramide (PerkinElmer) were used.

Furthermore, the RNAiso Plus, PrimeScript™ Reverse Transcriptase, Recombinant RNase Inhibitor, TBGreen. Premix Ex Taq™ (Tli RNaseH Plus), RR420 (Takara Bio Inc., Kyoto, Japan), RNeasy MinElute Cleanup Kit (QIAGEN, Tokyo, Japan), dNTP Mix and Oligo (dT)15 Primer (Promega, Tokyo, Japan), Low Input Quick Amp Labeling Kit, RNA6000 Nano Kit, Agilent Whole Human Genome DNA Microarray 4×44K v2, and Agilent Gene Expression Hybridization Kit (Agilent Technologies, Santa Clara, CA, USA) were also used in this study.

### 2.2. Preparation of sugi-pollen suspension and sugi-basic protein (SBP)

Japanese cedar (sugi) pollen (Biostir Inc., Osaka, Japan) was treated with ethanol, followed by preparation of a suspension with saline at a final concentration of 50 mg/mL. The obtained Sugi-pollen suspension was used for sensitization as a Sugi-pollen antigen to prepare a mouse model of pollinosis. Sugi-basic protein extract (SBP) was prepared by extracting the sugi pollen suspension with 0.9% NaCl containing 3% glycerol to yield a final concentration of 20 mg/mL Cryj-1 antigen. A quantitative assay for Cryj1 was performed using an LBIS Cryj-1 ELISA Kit (FUJIFILM Wako Pure Chemical Corporation, Osaka, Japan).

### 2.3. Preparation of soluble beta-glucan (SBG)

Crude SBG was obtained from culture supernatants of *Aureobasidium pullulans* (ATCC No. 2052). Purified SBG was prepared from the crude SBG by precipitation with 70% ethanol, followed by freeze-drying and dissolution in a sterile 0.9% NaCl solution. The amount the SBG was assayed according to the POPOD procedure using the β-glucan Assay Kit (Neogen Japan, Yokohama, Japan). In this study, we used 0.1% (1 mg/ml) β-glucan solution.

### 2.4. Animals

Female BALB/c mice were used in both studies, focusing on the pollinosis model and gut microbiota. As illustrated in Fig. 1A, the pollinosis model study comprised three groups: a control group (Ctr) that received PBS for 7 weeks, a pollinosis group (Polli) that was orally administered PBS for 7 weeks and sensitized twice with a sugi-pollen suspension, and an SBG-administered pollinosis group (SBG+Polli) that received SBG for 7 weeks and underwent the same sensitization with the sugi-pollen suspension. Figure 1B outlines the timeline and examinations for the microbiota study, which involved two animal groups: a control group (Ctr) and SBG group, which were orally administered PBS and SBG for 4 weeks, respectively. In both studies, each group consisted of seven animals for pollinosis study and six animals for microbiota examination.

**Figure 1.**
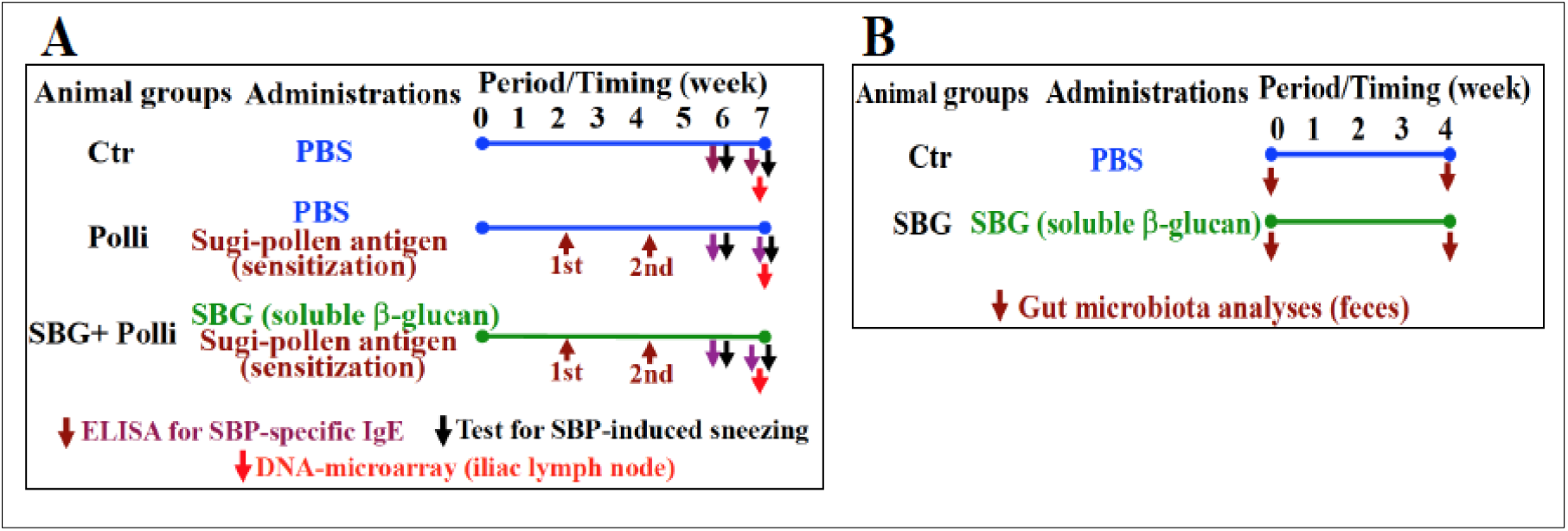
Outline for pollinosis examinations in mice and gut microbiota analyses. <u>**A**</u> shows pollinosis examinations. The control group (Ctr) and both the pollinosis group (Polli) and the SBG-administered pollinosis group (SBG+Polli) were orally given PBS and SBG for 7 weeks, respectively. Polli and SBC+Polli groups were sensitized twice, 1st at 2 weeks and 2nd at 4 weeks, with sugi pollen antigen. ELISA for SBP-specific IgE and a test for SBP-induced sneezing were performed at both 4 and 6 weeks. DNA microarray was carried out at 6 weeks. <u>**B**</u> displays gut microbiota analyses using stool samples from two mouse groups, the control group (Ctr) and the SBG-administered group (SBG). Ctr and SBG were administered with PBS and SBG for 4 weeks, individually.

### 2.5. Sensitization with sugi-pollen antigen

Mice of the Polli and SBC+Polli groups were sensitized twice: first at 2 weeks and then at 4 weeks. This was performed using intramuscular injections of a sugi-pollen suspension (100 mL/mouse) at the right and left bases of the tail without any adjuvant (see Fig. 1A). The sensitization procedure was conducted under ketamine hydrochloride anesthesia.

### 2.6. Preparation of Anti-SBP IgE Standard for ELISA

In preliminary studies, mice were sensitized with a sugi-pollen suspension, as mentioned above, to obtain anti-SBP sera for standard stock. Aliquots of the stocks were stored at −70°C until further use. The standard stock was serially diluted two-fold from 1:100 to 1:6400 to obtain an standard preparation with optical density from 1.8 to 0.1 at 450nm using the enzyme-linked immunosorbent assay (ELISA). Arbitrary units were assigned to the standards for stable titration, with the least diluted standard (1:100) being 100 and the most diluted standard (1:6400) being 2.0. The correlation between the dilution and units was always >0.9.

### 2.6. ELISA for SBP-specific IgE

Blood samples were collected from the tail vein of each mouse at 6 and 7 weeks of the examination period (Fig.1A) using a hematocrit capillary tube pretreated with heparin and then centrifuged at 1000 × g for 10 min to obtain plasma. Plasma levels of SBP-specific IgE were analyzed using a highly sensitive ELISA with the biotinyltyramide amplification system. Briefly, SBP (2μg/mL in PBS) was coated onto Immuno Module Maxisorp 96-well plates 4 °C overnight. After blocking with 1% casein Na at 4 °C overnight, standards and plasma samples, diluted 1:300 in 1% casein-Na prepared in PBS. Next, biotin-conjugated rat monoclonal anti-mouse IgE prepared at 0.8 μg/mL in 1% casein Na -PBS was reacted followed by successive application of streptavidin-horse radish peroxidase conjugate, biotynyltyramide, and streptavidin-horseradish peroxidase. Color was developed by adding a colorimetric substrate (TMB). The enzyme reaction was terminated with 2M sulfuric acid and the plate was read at 450 nm by subtracting the reference determined at 630 nm using an iMark microplate reader (Bio-Rad Laboratories Inc., Hercules, CA, USA). The plates were washed with 0.05% Tween-20 in PBS between each step except when sulfuric acid was added. The antibody titers of the samples were calculated using a calibration curve generated using a serially diluted standard.

### 2.7. Test for SBP-induced sneezing

The mice were individually placed in observation cages, and their sneezing behavior was monitored using a video camera at 6 and 7 weeks of the examination period (Fig. 1A). Both nasal cavities were pretreated with 10 mL/site of N-acetylcysteine at a final concentration of 3 mg/mL to perturb the mucin layer [14]. After washing twice with PBS (10 mL/site of PBS), the mice were intranasally challenged with 10 mL/site of an allergen, SBP. For each observation session, the number of sneezes during the 5 min after intranasal challenge with SBP was counted by at least three investigators. The average number of sneezes per session was calculated. In cases where the coefficient of variation exceeded 10%, the sneezing behavior of the mouse was re-examined by viewing the recorded video again.

### 2.8. Gut microbiota analyses

Fresh stool samples were collected in the morning at 0 and 4 weeks during the experimental periods (Fib.1B) and were stored at temperatures below −40 °C until analysis. DNA was extracted from 100–200 mg of frozen stool using a QIAamp DNA Stool Mini Kit (Qiagen, Valencia, CA, USA), following the manufacturer’s instructions. The concentration and purity of the extracted DNA were assessed by measuring the absorbance at 260 and 280 nm, respectively. Real-time PCR was conducted to quantify the 16S rRNA gene in 11 bacterial species/groups using a SYBR Green detection system. This was performed using the Mx3000P QPCR System (Agilent Technologies Inc., Santa Clara, CA, USA) using a SYBR Premix Ex Taq II (Tli RNase H Plus) Kit (Takara Bio Inc. Kyoto, Japasn) and specific primers for the target bacteria. The sequences of the primers (F: forward 5′ to 3′, R: reverse 5′ to 3′) are available upon request. The abbreviations for the six target bacteria are as follows: Lactobacilli (Lacto), Bifidobacteria (Bif), Clostridium butyricum (Cbu), Clostridium clostridiiforme (Ccl), Clostridium perfringens (Cpe), and Desulfovibrio (Dsv). A universal primer (Uni) set was employed to amplify the conserved regions of the 16S rRNA gene in bacteria and was used as a reference to express the relative quantity of the target bacteria, as detailed below.

The PCR reaction was conducted in a total volume of 10 μL, comprising 5 μL of SYBR Premix Ex Taq, 0.2 μL of ROX Reference Dye, 0.1 μM of each primer, and 25 ng of template DNA. The PCR conditions included an initial denaturation at 95 °C for 10 seconds, followed by 35 cycles of denaturation at 95 °C for 5 seconds and annealing/extension at 60 °C for 30 seconds. After each amplification, a dissociation curve analysis was performed to confirm PCR specificity. DNA standards for calibration were prepared from serially diluted PCR products that were previously amplified from macaque stool DNA using Uni primers. To quantify the target bacterial DNA in stool samples, the amount of the target DNA was calculated by dividing the quantity of interest by that obtained using Uni for normalization. Amplification and detection were performed in duplicate.

### 2.9. RNA isolation

To isolate RNA for DNA microarray analyses, iliac lymph node samples were collected within 3 h on the same day to minimize the influence of circadian clock genes. The samples were homogenized using RNAiso Plus. Total RNA was extracted as described previously [15, 16]. Briefly, RNA was treated with DNase using QIAGEN’s reagent in the aqueous phase and subsequently purified using the RNeasy MinElute Cleanup Kit from QIAGEN, following the manufacturer’s instructions. The quantity of RNA was assessed spectrophotometrically at 230, 260, 280, and 320 nm using an Ultrospec 2000 spectrometer (GE Healthcare Biosciences AB, Uppsala, Sweden). The RNA integrity number (RIN) of the DNA microarrays was determined using an Agilent 2100 Bioanalyzer (Agilent Technologies, Japan Ltd.). Only RNA samples of exceptional quality were used for the microarray examination, characterized by an A260/A230 ratio of 1.5, an A260/A280 ratio of 1.8, and an RIN value of 6.0.

### 2.10. DNA microarray

DNA microarray analysis was performed on the control group (Ctr) and two experimental groups, Polli and SBG + Polli. After synthesizing complementary DNA (cDNA), Cy3-labeled complementary RNA (cRNA) was generated and purified using a Low Input Quick Amp Labeling Kit following the manufacturer’s instructions. Reverse transcription was performed using a T7 promoter-oligo(dT) primer. The absorbance was measured at 260, 280, 320, and 550 nm to confirm that the labeled cRNA exhibited a concentration of > 6 pmol/mg of Cy3-CTP. Thereafter, the labeled cRNA was fragmented using the Gene Expression Hybridization Kit and applied to Whole Human Genome DNA Microarray 4°ø 44 K v2 slides. Following hybridization at 65 °C for 17 hours, the slides were washed with Gene Expression Wash Buffers 1 and 2 in accordance with the manufacturer’s instructions. The slides were subsequently scanned using a GenePix 4000 B scanner (Molecular Devices, San Jose, CA, USA). The scanned images were converted into a digital format and standardized using GenePix Pro 6.0 software (Molecular Devices).

### 2.11. Bioinformatic for Microarray Data Analyses

In the experimental group, we identified genes that exhibited > 2-fold upregulation or < 0.5-fold downregulation in expression compared with the control group. These genes were annotated, and databases such as PubMed, NCBI, and Google, and other information sources were used to search for references. We analyzed the immune-related functions of the annotated genes to gain insights into the potential mechanisms underlying the mitigating effects of SBG, while also utilizing additional relevant information sources for reference.

## 3. Results

In preliminary examinations of the dose-response of SBG within the range of 3–10 mg/kg in mice, we obtained reasonable results by oral administration of 10 mg/kg. We performed this study by using the dosage, 10 mg/kg, of SBG for oral administration in mice. Using the procedures shown in Fig.1(A and B), we obtained the following results.

### 3.1. Inhibitory effect of SBG on antigen-specific IgE production in pollinosiss mice model sensitized with sugi-pollen antigen

Figure 2A shows the plasma levels of SBP-specific IgE in pollinosis mice sensitized with Sugi-pollen antigen (Polli) compared with those of Ctr and SBG+Polli. The plasma levels of SBP-specific IgE in Polli were 8.4 and 12.2 units, respectively, at 6 and 7 weeks, respectively. In contrast, SBP-specific IgE levels in the SBG + Polli group decreased to 3.2 and 3.8 units, respectively, at 6 and 7 weeks. This indicated that pre-administration of SBG reduced SBP-specific IgE production in pollinosis mice (Polli). Thus, SBG inhibited SBP-specific IgE production in pollinated animals by 68–77% (Fig.2 B).

**Figure 2.**
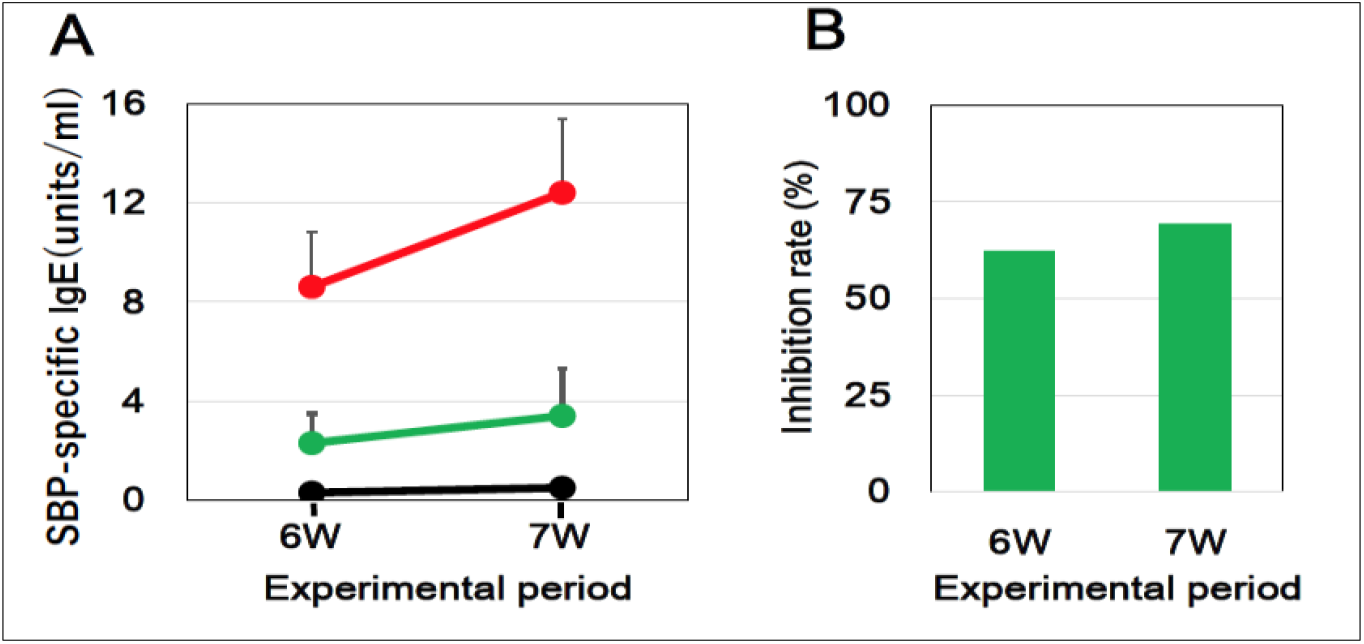
Inhibitory effect of SBG on SBP-specific IgE production in pollinosis mice. <u>**A**</u> shows plasma level of SBP-specific IgE in tree groups (Ctr, Polli, and SBG+Polli). SBP-specific IgE was measured By ELIS using plasma samples from the three groups at 6 and 7 weeks, respectively. <u>**B**</u> displays SBG-mediated inhibition to the SBP-specific IgE production in pollinosis mice (Polli) at 6 and 7weeks, respectively. Inhibition rate (%) was calculated from the plasma level of SBP-specific IgE in Polli and SBG+Polli.

### 3.2. Inhibitory effect of SBG on SBP-induced sneezing in pollinosis mice model

Figure 3A displays the number of SBP-induced sneezes among three groups, PBS, Polli, and SBG + Polli. The number of SBP-induced sneezes in Polli was 42 and 47 times/5 min, at 6 and 7 weeks, respectively. In contrast, the sneezing numbers in the SBG + Polli group were 14 and 18 times/min at 6 and 7 weeks, respectively. These results suggest that SBG administration inhibited SBP-induced sneezing in pollinosis animals by 64–72% (Fig.3 B).

**Figure 3.**
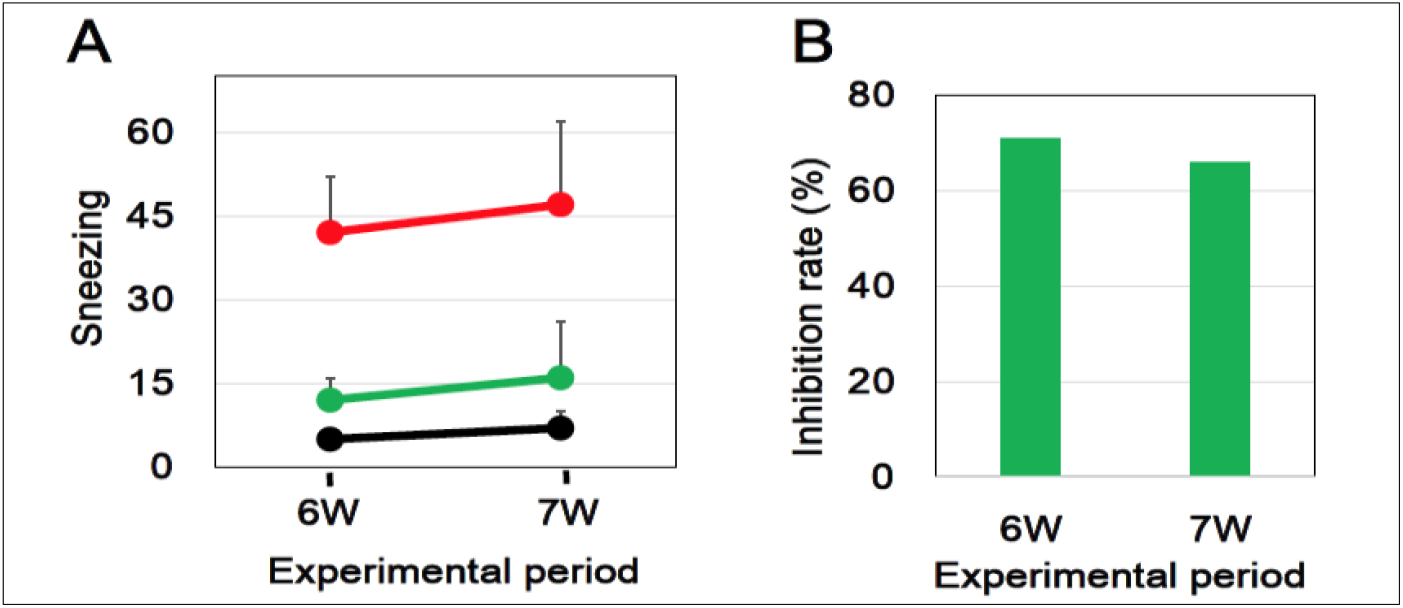
Inhibitory effect of SBG on SBP-induced sneezing in pollinosis mice. <u>**A**</u> shows the number (sneezing time /5 min) of SBP-induced sneezes among three groups, PBS, Polli, and SBG + Polli. <u>**B**</u> displays SBG-mediated inhibition to the SBP-induced sneezing in pollinosis mice (Polli) at 6 and 7weeks, respectively. Inhibition rate (%) was calculated from the sneezing time/5 min in Polli and SBG+Polli.

### 3.3. DNA microarray analyses and bioinformatics

To elucidate the mechanism underlying SBG-mediated inhibition of SBP-specific IgE production in pollinosis mice model, we performed DNA microarray analyses using RNA extracted from the iliac lymph nodes of three groups, Ctr, Polli, and SBG +Polli, at 7weeks.

#### 3.3.1. Amelioratively downregulated SBG-related genes

We compared the gene expression profiles of the two groups, Ctr and Polli, and identified 45 genes that were upregulated by more than 2.0-fold in Polli (data not shown). In the subsequent comparison of the upregulated gene expression profiles between Polli and SBG + Polli, we aimed to identify genes associated with the immune response, inflammation, and tissue repair that were downregulated in SBG + Polli. As shown in Table 1, we identified seven genes that were downregulated by more than 0.5-fold in the SBG + Polli and were related to the ameliorative action of SBG on pollinosis. These seven genes were referred to as amelioratively downregulated SBG-related genes. Table 1 lists the amelioratively downregulated SBG-related genes, including *Mnd1* (meiotic nuclear divisions 1 homolog), *Igk* (immunoglobulin kappa chain), *Igh* (immunoglobulin heavy chain variable region), *Aicda* (activation-induced cytidine deaminase), *S1pr2* (Sphingosine-1-phosphate receptor 2), *U76382*.*1* (immunoglobulin light chain mRNA), and *EndoU* (evolutionarily conserved single-stranded RNA binding protein, a novel regulator of B-cell activation-induced cell death).

**Table 1.** Amelioratively downregulated SBG-related genes.

| Gene symbol | Product: Description [Reference] | Fold change |  |
| --- | --- | --- | --- |
|  |  | Polli<sup>***</sup> / Ctr<sup>*</sup> | SBG+Polli<sup>***</sup> / Polli<sup>***</sup> (SBG+Polli<sup>***</sup> / Ctr<sup>*</sup>) |
| *Mnd1* | MIND1: Meiotic nuclear divisions 1 homolog (S. cerevisiae) [22] | 3.6 | 0.3<br>(1.2) |
| *Igk* | IgK: Immunoglobulin kappa chain [19- 21] | 6.0 | 0.3<br>(2.0) |
| *Igh* | IgH: Immunoglobulin heavy chain variable region [19- 21] | 6.3 | 0.3<br>(2.1) |
| *Aicda* | AICDA: Activation-induced cytidine deaminase [23] | 25.4 | 0.4<br>(10.2) |
| *S1pr2* | S1PR2: Sphingosine-1-phosphate receptor 2 [24] | 2.8 | 0.4<br>(1.1) |
| *U76382.1* | Immunoglobulin kappa light chain, Anti-CD4 immunoglobulin light chain mRNA [19- 21] | 2.2 | 0.5<br>(1.1) |
| *Endou* | EndoU: Endonuclease, Poly(U) Specific, Novel regulator of B-cell activation-induced cell death [25] | 7.6 | 0.5<br>(3.8) |

<sup>\*</sup>Ctr: Control group, <sup>\*\*</sup>Polli: Sugi-pollen sensitized group, <sup>\*\*\*</sup>SBG+Polli: SBG pre-administrated Polli group.

#### 3.3.2. Amelioratively upregulated SBG-related genes

By comparing the gene expression profiles between the two groups, Ctr and Polli, we identified 73 genes that were downregulated by a factor of 0.5 or lower in the Polli group (data not shown). We then analyzed the downregulated gene expression profiles between Polli and SBG + Polli groups to select genes associated with immune response, inflammation, or tissue repair that were upregulated in the SBG + Polli group. As shown in Table 2, we extracted twelve genes that were upregulated by a factor of 2.0 or greater in the SBG + Polli. These genes appear to be associated with SBG-mediated amelioration of pollinosis. These 12 genes are referred to as amelioratively upregulated SBG-related genes. Table 2 lists these amelioratively upregulated genes, which include *Ngfr* (Nerve growth factor receptor), *Prg2* (Proteoglycan 2, bone marrow), *Ear6* (Eosinophil-associated, ribonuclease A family, member 6), *Chi3l3* (Chitinase 3-like 3), *Pon1* (Paraoxonase 1), *Cd209b* (CD209b antigen, transcript variant 2), *Ear1* (Eosinophil-associated, ribonuclease A family, member 1), *Ms4a8a* (Membrane-spanning 4-domains, subfamily A, member 8A), *Camp* (Cathelicidin antimicrobial peptide), *Ltf* (Lactotransferrin), *Elane* (Elastase, neutrophil expressed), and *Ngp* (Neutrophilic granule protein).

**Table 2.** Amelioratively upregulated SBG-related genes.

| Gene Symbol | Product: Description [Reference] | Fold Change |  |
| --- | --- | --- | --- |
|  |  | Polli<sup>***</sup> / Ctr<sup>*</sup> | SBG+Polli<sup>***</sup> / Polli<sup>***</sup> (SBG+Polli<sup>***</sup> / Ctr<sup>*</sup>) |
| *Ngfr* | NGFR: Nerve growth factor receptor (TNFR superfamily, member 16) [26] | 0.1 | 3.0<br>(0.3) |
| *Prg2/Mbp* | PRG2/MBP: Proteoglycan 2 bone marrow, also named eosinophil major basic protein (MBP) [27] | 0.1 | 3.0<br>(0.3) |
| *Ear6* | EAR6: Eosinophil-associated, ribonuclease A family, member 6 [28] | 0.1 | 3.0<br>(0.3) |
| *Chi3l3/Ym1* | CHI3L3/YM1: Chitinase 3-like 3, also named Ym1 [29,30,31] | 0.2 | 2.5<br>(0.5) |
|  |  | Polli<sup>**</sup> / Ctr<sup>*</sup> | SBG+Polli<sup>***</sup> / Polli<sup>**</sup><br>(SBG+Polli<sup>***</sup> / Ctr<sup>*</sup>) |
| *Ngfr* | NGFR: Nerve growth factor receptor (TNFR superfamily, member 16) [26] | 0.1 | 3.0<br>(0.3) |
| *Prg2/Mbp* | PRG2/MBP: Proteoglycan 2 bone marrow, also named eosinophil major basic protein (MBP) [27] | 0.1 | 3.0<br>(0.3) |
| *Ear6* | EAR6: Eosinophil-associated, ribonuclease A family, member 6 [28] | 0.1 | 3.0<br>(0.3) |
| *Chi3l3/Ym1* | CHI3L3/YM1: Chitinase 3-like 3, also named Ym1[29,30,31] | 0.2 | 2.5<br>(0.5) |
| *Pon1* | PON1: Paraoxonase 1 [32] | 0.2 | 3.0<br>(0.6) |
| *Cd209b* | CD209b antigen, transcript variant 2 [33] | 0.2 | 2.5<br>(0.5) |
| *Ear1* | EAR1: Eosinophil-associated, ribonuclease A family, member 1 [34] | 0.2 | 2.5<br>(0.5) |
| *Ms4a8a* | MS4A8A: Membrane-spanning 4-domains, subfamily A, member 8A [35] | 0.3 | 2.3<br>(0.7) |
| *Camp* | CAMP: Cathelicidin antimicrobial peptide [36] | 0.3 | 2.3<br>(0.7) |
| *Ltf* | LTF: Lactotransferrin [37] | 0.4 | 2.0<br>(0.8) |
| *Elane* | ELANE/NE: Elastase, neutrophil expressed [38] | 0.4 | 2.0<br>(0.8) |
| *Ngp* | NGP: Neutrophilic granule protein [41] | 0.5 | 2.0<br>(1.0) |

<sup>\*</sup>Ctr: Control group, <sup>\*\*</sup>Polli: Sugi-pollen sensitized group, <sup>\*\*\*</sup>SBG+Polli: SBG pre-administrated Polli group.

### 3.4. Effect of SBG on gut microbiota

We also examined the effect of oral SBG administration on the gut microbiota of mice in a separate study, as outlined in Fig. 1B. Because of significant individual differences in the gut microbiota, conducting comparative studies on changes between the control and SBG-administered groups, which consisted of a limited number of animals (i.e., 6–8 mice), was challenging. Therefore, in this study, we utilized stool samples collected from the same animals before (W0) and after (W4) SBG administration to compare the effects of SBG on gut microbiota at these two time points, W0 and W4. We performed real-time PCR using DNA extracted from stool samples to assess the levels of six target bacteria, which were beneficial bacteria including Bifidobacteria (Bif); Clostridium butyricum (Cbu); and Lactobacilli (Lact) and harmful bacteria including C. clostridiiforme (Ccl); C. perfringens (Cpe); and Desulfovibrio (Dsv).

As shown in Figure 4A, the levels of three beneficial bacteria, Bif, Cbu, and Lact, increased 8.7-, 1.5-, and 2.4 -fold, respectively, after four weeks of SBG administration. Conversely, the levels of three harmful bacteria, Ccl, Dsv, and Cpe, barely changed or decreased by 0.4-fold during four weeks of SBG administration (Fig. 4B). Very little alteration in the levels of both beneficial and dangerous bacteria was noted in fecal samples from control mice following 4 weeks of PBS administration. (data not shown). These results indicated that SBG administration led to an increase in beneficial bacteria, particularly Bif, in the gut microbiota.

**Figure 4.**
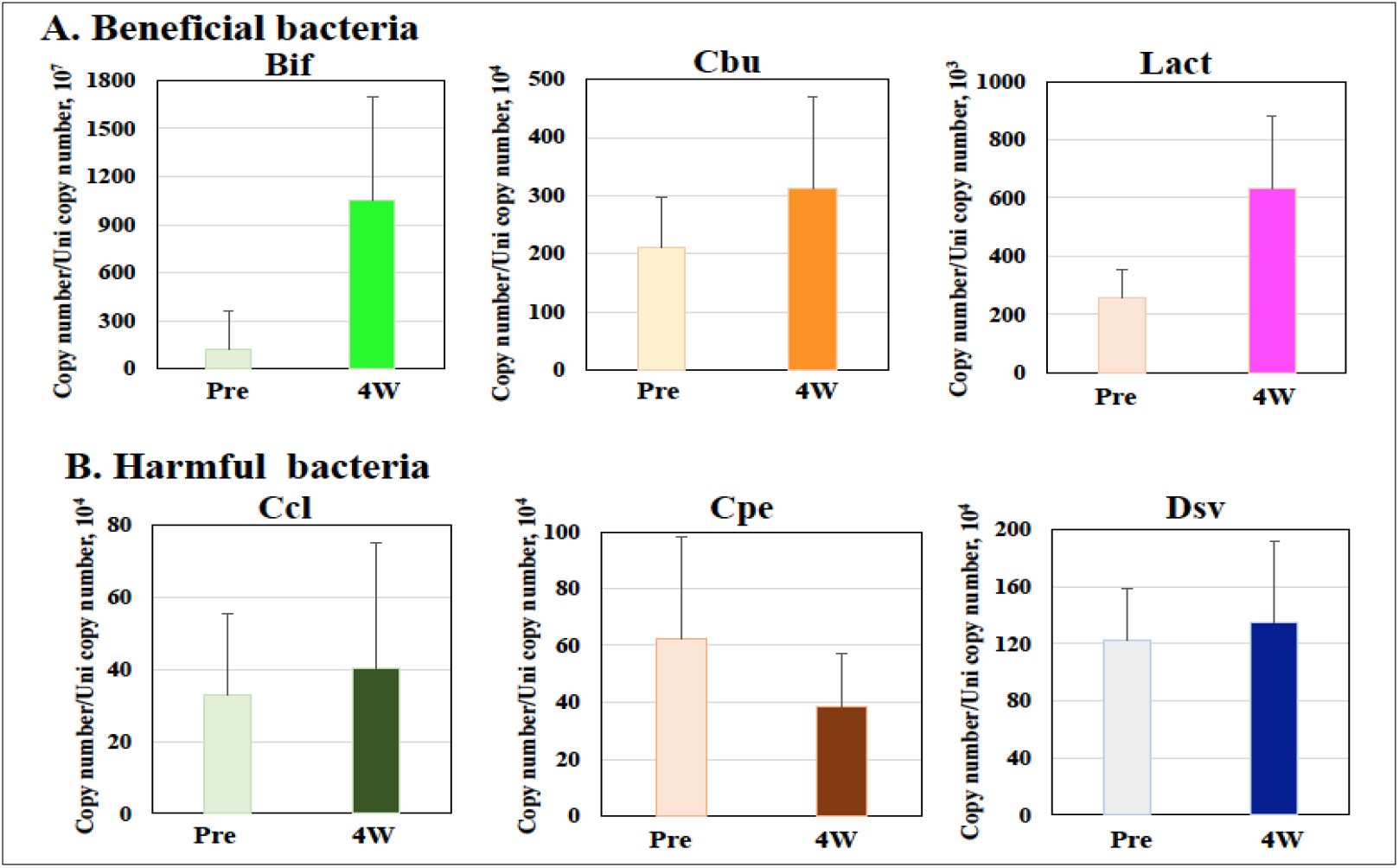
Effect of SBG on gut microbiota. <u>**A**</u> shows the levels of three beneficial bacteria, Bifidobacteria (Bif); Clostridium butyricum (Cbu); and Lactobacilli (Lact), increased 8.7; 1.5; and 2.4-fold, respectively, after four weeks of SBG administration. <u>**B**</u> displays the levels of three harmful bacteria, C. clostridiiforme (Ccl); C. perfringens (Cpe); and Desulfovibrio (Dsv), barely changed (Ccl and Dsv) or decreased by 0.5-fold (Cpe) during four weeks of SBG administration

## 4. Discussions

### 4.1. Sugi-pollinosis, IBGs, and SBGs

Sugi-pollinosis is caused by the SBP-specific IgE antibodies, which bind to FcεRI, a high-affinity receptor that is expressed on the surface of mast cells and/or basophils. The initial step in the development of pollinosis includes production of SBP-specific IgE antibodies.

Therefore, certain foods and related materials that suppress the production of SBP-specific IgE may help alleviate sugi-pollinosis.

There are two types of β-glucans: IBGs and SBGs. IBGs activate dectin-1 signaling in various immune-inflammatory cells, including B and T cells, antigen-presenting cells, mast cells, and granulocytes (neutrophils and eosinophils). Activation of these cells by IBGs begins with their binding to dectin-1, which is present on the cell surface. This triggers oligomerization and activates the Src-Syk signaling cascade, leading to an immune-inflammatory response [1–4]. In contrast, SBG acts as a dectin-1 antagonist; it binds to dectin-1 but does not initiate any intracellular signaling pathway [1]. Consequently, SBGs inhibit the action of IBGs by competing for binding to dectin-1 and preventing IBG-mediated activation of the intracellular machinery [12,13].

During the immune response, IBGs function as adjuvants leading to the production of specific antibodies, including IgE. Pollen grains contain allergy-related components, allergens, and IBGs, similar to adjuvants, which initiate allergic immune responses [8]. Japanese cedar pollen contains IBG polysaccharides and exhibits IgE-inducing activity in mice [9]. Kanno et al. reported that sugi pollen contains IBGs that stimulate dectin-1 signaling and demonstrates a potent adjuvant effect under allergic conditions, promoting Th2 imbalance and SBP-specific IgE production [11]. In the present study, we leveraged the adjuvant effect of IBGs to successfully establish a sugi-pollinosis mouse model through sensitization with only sugi pollen. This was done without the use of other adjuvants such as alum, which is commonly used to create allergy models associated with specific IgE production.

Previous studies have reported that SBGs exhibit anti-allergic effect [12, 13], including in humans [14]. We confirmed the anti-allergic actions of SBGs by demonstrating the inhibition of both SBP-specific IgE production and SBP-induced sneezing after oral pre-administration in a sugi-pollinosis mouse model (Figs. 2 and 3). The anti-allergic effect mediated by SBGs likely results from their competition with adjuvant IBGs for binding to dectin-1, which subsequently inhibits intracellular mechanisms involved in allergic immune responses. However, detailed information regarding the molecular mechanisms underlying the anti-allergic action of orally administered SBGs has not yet been adequately gathered.

### 4.2. DNA microarray analyses and Iliac lymph nodes

An efficient method for studying the expression levels of most genes in an organism’s functional genome, particularly the transcriptional product (mRNA), is through DNA microarray techniques, which provide high-throughput analysis of transcripts/mRNAs. In our previous studies on sublingual vaccination [15,16], we elucidated novel molecular events within the immune system and their related responses using a DNA microarray approach. In the present study, we further studied the mechanism underlying the anti-allergic action of orally administered SBGs in a model of sugi-pollen allergy model. DNA microarray analyses were conducted using RNA extracted from the iliac lymph nodes, as these nodes are local lymphatic tissues located near the tail base muscle, where the sugi-pollen antigen was intramuscularly injected to sensitize the animals to pollinosis. A highly effective immune response, known as sensitization, has been reported to occur in lymphocytes from the iliac lymph nodes following the intramuscular injection of a sensitizing antigen at the tail base in mice [17]. Therefore, a method involving the iliac lymph nodes, through intramuscular sensitization at the tail base and subsequent collection of these lymph nodes, was established to produce specific antibodies [18]. This method has also been utilized in studies examining allergies and related immune-inflammatory responses.

### 4.3. Amelioratively downregulated SBG-related genes

Using DNA microarray analysis, we identified four additional SBG-related genes that were downregulated, including *Mnd1* (meiotic nuclear division 1 homolog), *Aicda* (activation-induced cytidine deaminase), *S1pr2* (Sphingosine-1-phosphate receptor 2), and *EndoU* (evolutionarily conserved single-stranded RNA-binding protein, a novel regulator of B-cell activation-induced cell death) (Table 1). MND1 is involved in DNA repair during cell growth and is related to the cell cycle, proliferation, invasion, and migration of cells within the lymph node microenvironment [22]. Consequently, MND1 appears to be associated with the stimulation of the immune response through the infiltration of immune cells into the lymph nodes. Reportedly, AICDA exhibits highest expression in the spleen of OVA-sensitized allergic mice [23], with its gene expression significantly promoted by IL-21 and IL-17A [23]. S1PR2 is a receptor of sphingosine-1-phosphate (SIP), an important signaling molecule that is associated with allergic inflammatory diseases, including pollen allergies, asthma, and anaphylaxis [24]. The interaction between S1PR2 and SIP on mast cells enhances their activation, contributing to allergic inflammation [24]. EndoU serves as a critical regulator of the pathway that controls B cell survival and antigen responsiveness, contributing to B cell tolerance [25]. EndoU is associated with the survival of pathogenic B cells, such as those responding to sugi-pollen antigens in sugi-pollen allergies. As these four genes directly or indirectly participate in the dysregulated stimulation of the immune system, the upregulation of these genes in Polli mice correlates with an allergic state in local lymphoid tissues, such as the iliac lymph nodes. Conversely, the downregulation of these upregulated genes in SBG-administered mice (SBG + Polli) ameliorated the dysregulated immune response, including allergic reactions. Therefore, SBG administration appeared to ameliorate sugi-pollen allergies.

### 4.4. Amelioratively upregulated SBG-related genes

As revealed by the ameliorative upregulation of SBG-related gene (Table2), we uncovered 12 genes whose expressions were upregulated in SBG-administrated mice, SBG + Polli, comparison with those in sugi-pollen allergy mice, Polli. NGFR is a nerve growth factor (NGF) receptor that dampens the inflammatory response and limits tissue damage through the NGFR-NGF interaction [26]. Thus, NGFR likely contributes to regulatory feedback on the development and maintenance of chronic inflammation, including allergies. PRG2, also known as eosinophil major basic protein (MBP), is involved in the antiparasitic defense system as a helminth toxin and in immune hypersensitivity reactions. MBP also participates in tissue remolding in allergic atopy through its pro-angiogenic effects [27]. EAR6 belongs to the eosinophil-derived RNase A superfamily and is involved in innate and Ag-specific immune responses to danger signals such as endogenous multifunctional immune alarmins [28]. CHI3L3/YM1 is primarily produced by macrophages and has multifunction [29]. CHI3L3/YM1 stimulates the Th2 response in the early phase of allergic inflammation [30], but also inhibits the Th2 response by reducing the gene expression of Th2 cytokines (IL-5 and IL-13), resulting in the regulation of Th2 imbalance [31]. Thus, upregulation of CHI3L3/YM1 in SBG+Polli is thought to relate to anti-allergic effect of SBG administration. PON1 is a major antioxidant enzyme and acts to ameliorate Th2-mediated inflammation by downregulating IL-4 and IL-13 and upregulating TGF-β [32]. EAR1 associates with anti-inflammatory reaction to regulate macrophage and polarization and block NF-κB activation [34]. MS4A8A is a CD20 homolog expressed in M2 macrophages and plays a role in integrating anti-inflammatory signals into M2-like macrophages [35]. CAMP is an immunomodulatory peptide mainly secreted by immune cells, macrophages and lymphocytes, and has a regulatory role of inflammatory mediators, such as TNFα [36]. LTF has multiple physiological functions, including anti-inflammatory effects, and exhibits protective effects by reducing the expression of Th2 cytokines and secretion of allergen-specific antibodies in an OVA-induced allergic mouse model [37]. ELANE/NE plays a role in dampening inflammatory reactions by degrading various pro-inflammatory cytokines, including IL-1, TNF, and IL-6 [38]. Neutrophils are known to participate in both the initiation and resolution of inflammation [39,40], and neutrophil granule proteins (NGP) play a role in minimizing inflammatory damage to neighboring tissues through modulation of neutrophil trafficking, migration, and/or extracellular trap formation [41]. Following pollen antigen sensitization in Poll mice, the iliac lymph nodes were enlarged due to inflammatory events accompanied by the infiltration of immune-inflammatory cells, macrophages, lymphocytes (B and T cells), and granulocytes (neutrophils, basophils, and eosinophils) (data not shown). In turn, SBG administration to SBG +Poll mice relieved iliac lymph node enlargement. As mentioned above, the 12 upregulated genes were expressed in immune-inflammatory cells within the iliac lymph nodes, and SBG administration dampened the inflammatory events by upregulating the expression of these genes in the lymph nodes.

### 4.5. Mechanism underlying anti-allergy action mediated by SBG on sugi pollinosis

Figure 5 illustrates potential molecular events occurring in the aforementioned immune-inflammatory cells of iliac lymph nodes, which are in an allergic state (L1 and L1-ii) induced by sugi pollen (SBP+IBG) or an ameliorated allergic condition (L2 and L2-ii) resulting from SBG. Figure 5 demonstrates that SBG inhibits the binding of IBG to dectin-1 on several immune-inflammatory cells, together with the modulation of upregulated expression of SBG-related genes linked to the IBG-mediated allergic responses in the lymph nodes (L1 and L2). This figure also explains the alleviation of IBG-stimulated immune-inflammatory reactions in the related cells (LI-ii and L2-ii).

**Figure 5.**
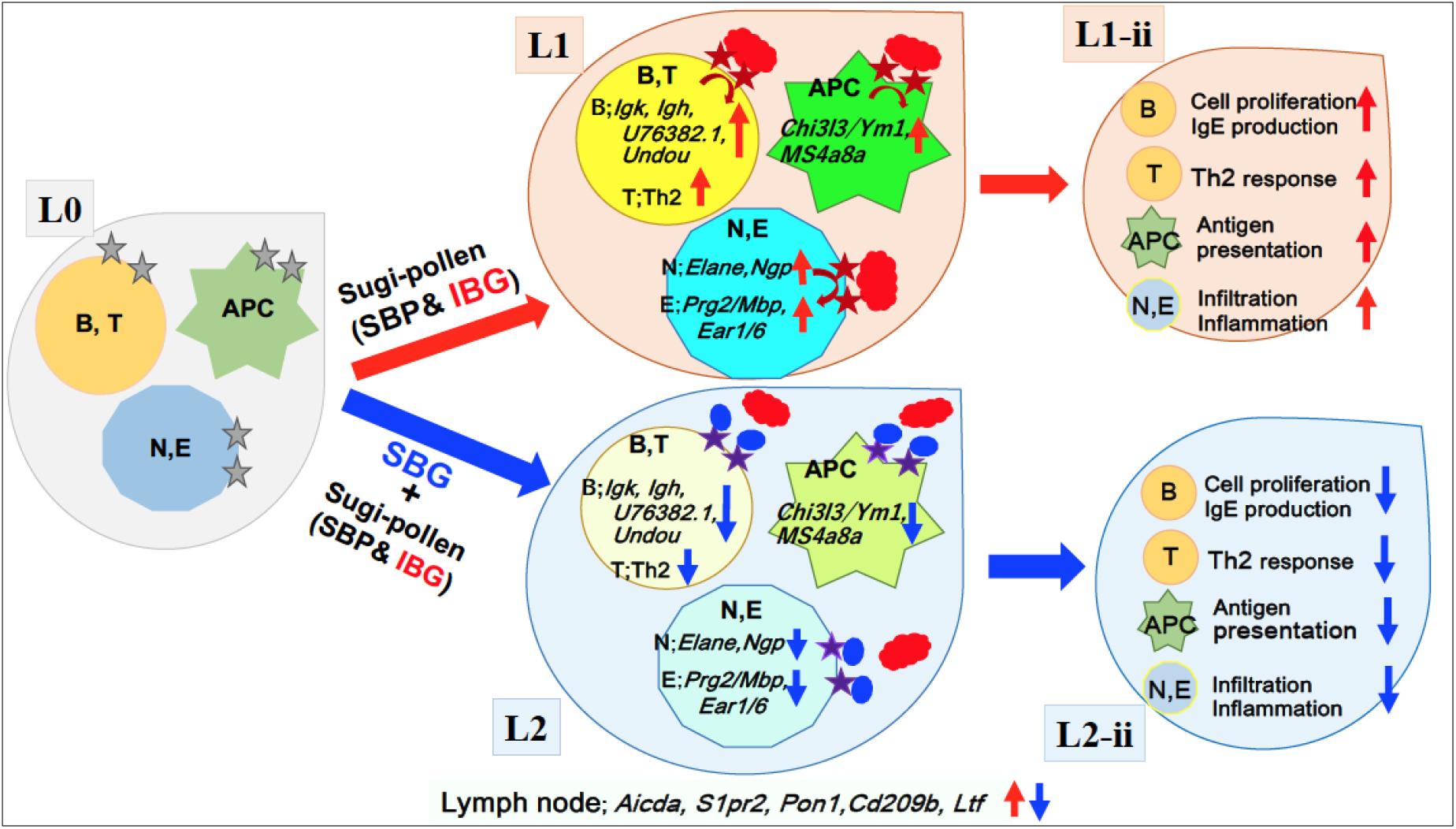
Possible mechanism underlying anti-allergy action mediated by soluble **β**-glucan (SBG) on Japanese cedar pollinosis. L0, L1/LI-ii, and L2/L2-ii indicate steady state, allergic condition generated by sugi pollen (SBP+IBG), and mitigated status caused by SBP, respectively, of iliac lymph nodes. B, T, APC, N, and E mean B- and T-cells, antigen-presenting cells, neutrophils, and eosinophils, respectively. Gray, brown, and purple stars represent the dectin-1 receptor expressed on the aforementioned cells, separately. Red solid clouds in both L1 and L2, along with a blue solid circle in L2, indicate insoluble β-glucan (IBG) and soluble β-glucan (SBG), individually. In L1, genes with red upward arrows indicate sugi pollen (SBP+IBG)-induced upregulated expression in the above-mentioned immune-inflammatory cells. In L2, genes with blue downward arrows mean amelioratively downregulated *SBG*-related ones in these cells. In L1-ii, immune-inflammatory reactions with red upward arrows indicate sugi pollen (SBP+IBG)-induced activation status in each cell. On the other hand, in L2-ii, these reactions with blue **d**ownward arrow**s** represent SBG-mediated alleviating states in the immune-inflammatory cells of iliac lymph nodes.

### 4.6. Suppression of mast cell activation by SBG

Mast cells play a pivotal role in initiating and sustaining inflammation, particularly in allergic reactions. Upon re-exposure to allergens, IgE-mediated aggregation of the IgE receptor, FcεRI, occurs on the mast cell surface, leading to rapid degranulation of the mast cells [42].

Antigen-induced sneezing is a common allergic symptom. This occurs when intranasal mast cells are activated by the binding of antigens to the antigen-specific IgE and FcεRI complex, which is followed by degranulation and the release of inflammatory mediators, including histamine. SBG directly inhibits antigen-induced mast cell activation, resulting in anti-allergic effects [43, 44]. In the current study, we also found that oral administration of SBG inhibited SBP-induced sneezing in SBG + Polli mice (Fig. 3A and B). This inhibition is believed to stem from the suppressive effect of SBG on mast cell activation via dectin-1. Mast cells are known to express dectin-1, and a dectin-1 antagonist, SBG (laminarin), has been shown to block the signaling pathways that lead to mast cell activation through dectin-1 signaling [45].

Another mechanism contributing to the SBG-mediated inhibition of mast cell activation has also been reported. In this instance, antigen-mediated mast cell activation is suppressed by short-chain fatty acids (SCFAs), which are metabolic products of the gut microbiota [46]. A previous study indicated that oral administration of yeast-derived SBG acted as a prebiotic, altering the gut microbiota to increase the abundance of beneficial bacteria, resulting in elevated levels of fecal SCFAs, including acetic, propionic, and butyric acids [47]. In this study, we also found that SBG administration over a 4-week period resulted in changes in the gut microbiota profile and an increase in beneficial bacteria, specifically Bif (Fig 4). Additionally, oral administration of *β*-glucan caused an increase in beneficial bacteria, accompanied by higher levels of SCFAs, including butyric acid, lactic acid, and propionic acid (48, 49). Although we did not obtain direct evidence for SBG-mediated accumulation of SCFAs in this study, it is plausible that SBG functions as a prebiotic, suppressing mast cell activation through SCFAs produced by gut microbiota, which may help mitigate sugi-pollenosis and other related allergies.

Figure 6 presents a potential mechanism by which SBG inhibits mast cell activation. Briefly, SBG competes with IBS for binding to dectin-1, reduces FcεRI stimulation due to lower levels of SBP-specific IgE, and is modulated by SCFA regulation. These factors contribute to the suppression of mast cell degranulation, leading to inhibit histamine release and reduced sneezing.

**Figure 6.**
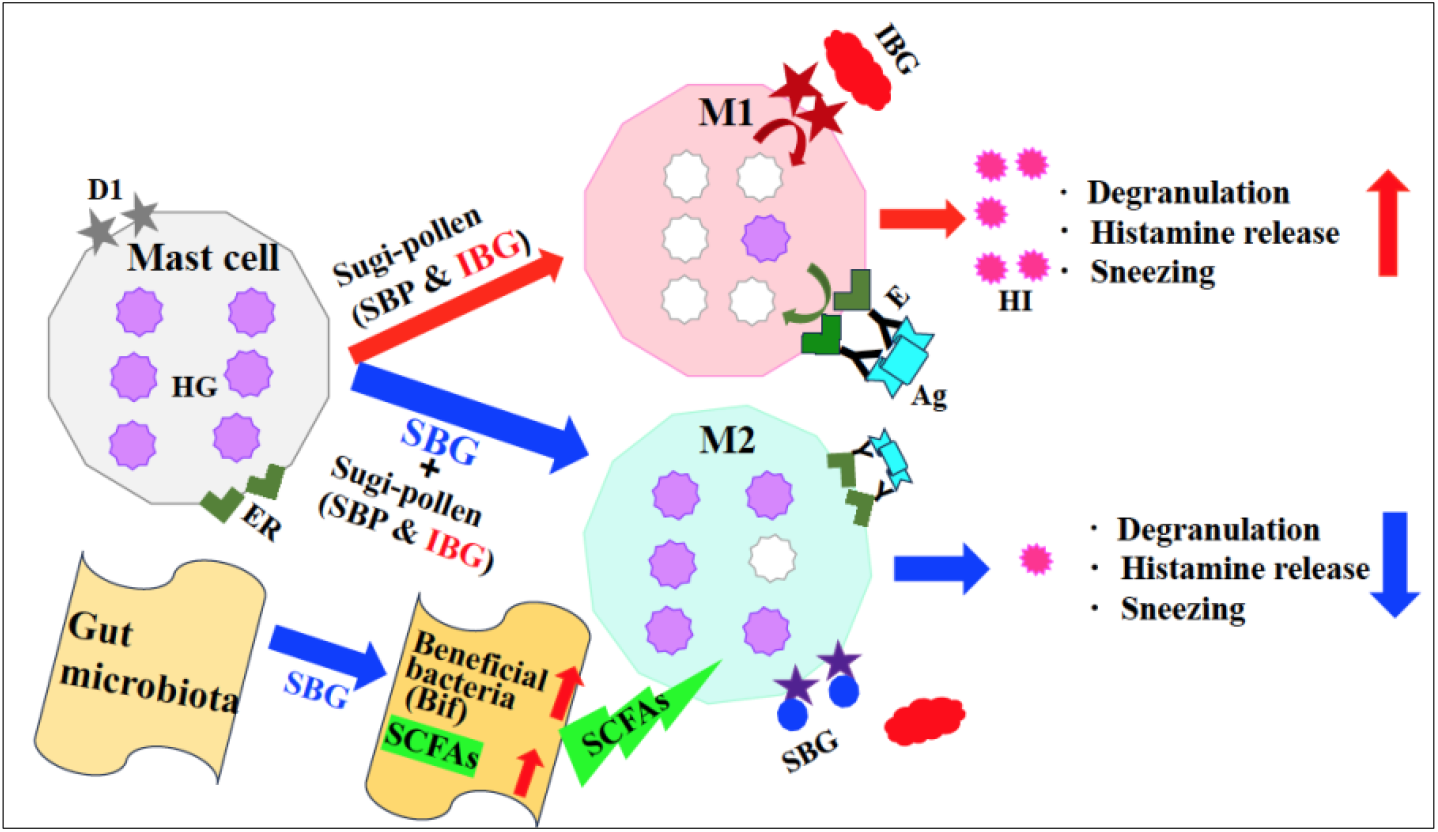
Potential mechanism for soluble **β**-glucan (SBG)-mediated inhibitory action on mast cell activation. M1and M2 mean mast cells of an activated/degranulated sate mediated by sugi pollen (M1) and an ameliorated status induced by SBG, respectively. D1, HG, ER, SCFAs, IBG, E, Ag, SBG, and HI mean dectin-1receptor, histamine granule, FcεRI, short-chain fatty acids, insoluble *β*-glucan particles, SBP-specific IgE, SBP antigen, soluble **β**-glucan, and histamine, individually.

## 5. Conclusion

IBG acted as a dectin-1 agonist and was present in sugi pollen as a polysaccharide adjuvant that enhanced the immune-inflammatory response. This response led to the production of SBP-specific IgE and the activation of mast cells, resulting in sneezing associated with SBP. In contrast, SBP, a dectin-1 antagonist derived from black yeast, mitigated sugi pollen allergy by reducing SBP-specific IgE production and decreasing SBP-induced sneezing in a mouse model of sugi pollinosis. The anti-allergic effects of SBP were mediated by the preventing of the IBS-stimulated immune-inflammatory response in B and T cells, antigen-presenting cells, and granulocytes. Additionally, SBP inhibited mast cell activation by reducing SBP-specific IgE levels and promoting beneficial bacteria, which subsequently leads to the production of short-chain fatty acids (SCFAs). Therefore, SBP and related functional foods and/or beverages appear to be advantageous for controlling allergies.

## Author Contributions

Conceptualization, M.T., T.S. and S.N.; Investigation, F.M. and S.N.; Methodology, F. M.; Interpretation of data and Project administration, M.T. and K.M.; Supervision, M.T. and T.S.; Validation, K.M.; Writing – original draft, M.T. and S. N.

## Institutional Review Board Statement

This study was conducted according to the guidelines of Institutional Animal Care and Committee Guide of Intelligence and Technology Lab, Inc. (ITL) based on the Guidelines for Proper Conduct of Animal Experiments and approved by the Animal Care Committee of the ITL (approved number: AE2023011 date: 11 September 2023). This study was also approved by the ITL Biosafety Committee (approved number: BS2023011, date: 11 September 2023).

## Informed Consent Statement

Not applicable.

## Data Availability Statement

Data are available from S.N. upon reasonable request.

## Acknowledgments

We thank to Kazuhiro Kawai, Masaki Matsuura, and Makiko Itoda for their invaluable technical assistances.

